# Using Ordinary Differential Equations to Explore Cancer-Immune Dynamics and Tumor Dormancy

**DOI:** 10.1101/049874

**Authors:** Kathleen P. Wilkie, Philip Hahnfeldt, Lynn Hlatky

## Abstract

Cancer is not solely a disease of the genome, but is a systemic disease that affects the host on many functional levels, including, and perhaps most notably, the function of the immune response, resulting in both tumor-promoting inflammation and tumor-inhibiting cytotoxic action. The dichotomous actions of the immune response induce significant variations in tumor growth dynamics that mathematical modeling can help to understand. Here we present a general method using ordinary differential equations (ODEs) to model and analyze cancer-immune interactions, and in particular, immune-induced tumor dormancy.

## I. INTRODUCTION TO CANCER-IMMUNE INTERACTIONS

Tumor growth within a host is defined by the interactions the cancer cells have with the local environment, including many functionally different immune cells. The intercellular signaling between cancer and stromal cells creates an ever-evolving milieu of cytokines that determine whether the immune cell function will be tumor-promoting or tumor-inhibiting [1]-[6] - and thus the overall tumor growth dynamics. Indeed, growth dynamics vary among lesions within a host [7], across hosts, and following treatments [8], depending on inherent variability in factors such as the sensitivity of cancer cells or the response of host cells to communication signals [9], [10].

Immune-induced tumor dormancy is a transient state of cancer progression during which the abnormal cells and microenvironment may evolve over time, but tumor mass remains constant [11]-[13]. Dormancy can occur at any stage and may terminate in either tumor elimination or regrowth [14]. In some instances, the dormancy may persist throughout the patient’s lifespan, resulting in an asymptomatic cancer [15]. Immunotherapy aims to boost the cytotoxic immune response to eliminate the disease, but can result in variable and non-intuitive response dynamics, including dormany or even increased tumor mass prior to regression [8].

An analogy to aid understanding of these complex tumor dynamics lies in the motion of a tumor mass rolling within an immune “potential well”, where escape to one side represents tumor elimination, escape to the other represents tumor progression, and the “trapped” motion within the well represents tumor dormancy [16], see Fig. 1. Understanding the dynamics of tumor growth within a patient is important for treatment planning, especially for secondary or tertiary treatments where environmental regulation is aberrant [9].

**Figure 1.**
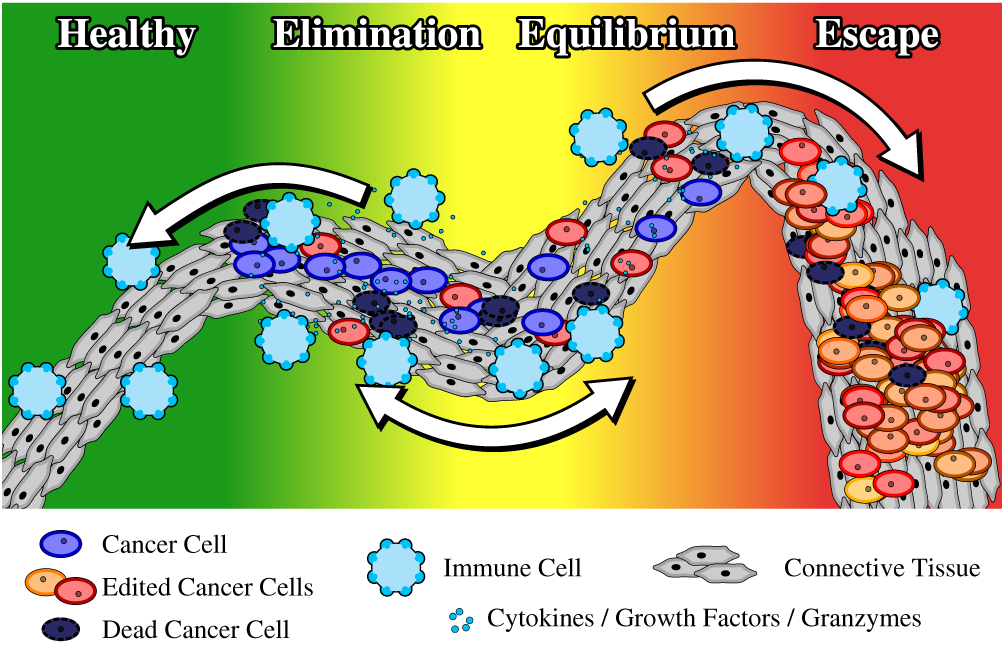
Visualization of cancer-immune dynamics as motion within a potential well (derived from Fig. 10.1 in [16]).

## II. RESULTS AND INSIGHTS FROM APPLYING THIS METHOD

With this model we demonstrate how tumor dormancy is a transient state that necessarily ends in either tumor elimination or escape [9], see Fig. 2. This result agrees with real world observations and contrasts the standard (mathematical) view of dormancy as a stable equilibrium [16]-[21] that persists for a long time. This model also captures known non-intuitive tumor dynamics such as accelerated repopulation [22]-[24] and anomalous periods of growth prior to regression that have been observed post tumor treatment [8].

**Figure 2.**
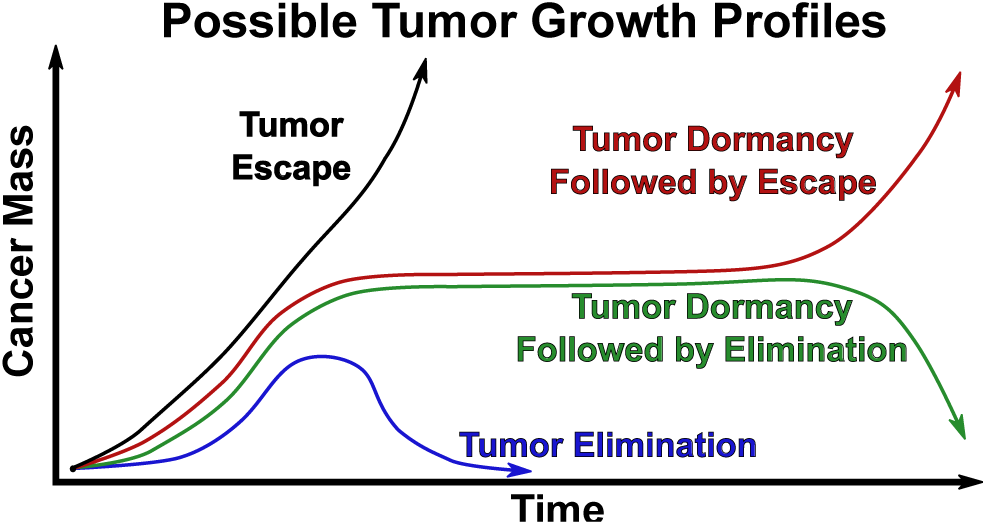
Possible tumor growth profiles including elimination, escape, and transient periods of tumor dormancy.

The model also demonstrates the effect of cell-level sensitivity to environmental regulation, showing how inherent variability of response to signals can explain disparate patient outcomes for similar treatment protocols [9], [10]. The method is proposed as a proxy to help analyze patient variability and treatment outcomes to improve understanding of why specific treatments work for some patients but not others.

## III. QUICK GUIDE TO THE METHODS

The method of applying ODEs to study cancer-immune dynamics requires, in the most basic form, prescribing equations for the rate of change over time of the cancer cell and immune cell populations, thus creating a system of two ODEs that are solved simultaneously. Several reviews can be found in the literature [16], [25].

In this approach [9] we do not model specific cytokine concentrations, but rather prescribe functional forms that encompass the net result of many cytokines on the bulk population. This allows for a minimally parameterized model that is easier to validate and more amenable to demonstrating the fundamental dynamics governing the cellular interactions.

Fig. 3 depicts the main mechanisms by which the cancer and immune cells interact. Each population has a net growth rate that is defined as the birth rate less the death rate for that population. Immune cells can promote cancer growth through inflammatory processes that increase blood flow and stimulate angiogenesis, bringing growth factors and nutrients to the tumor site [3]. They can also induce apoptosis through cytotoxic action [4], [26]. Cancer cells, on the other hand, can inactivate immune cells and develop mechanisms to evade immune detection [12], [27], [28]. They also recruit immune cells to the tumor by disrupting tissue architecture and producing chemokines and cytokines [3].

**Figure 3.**
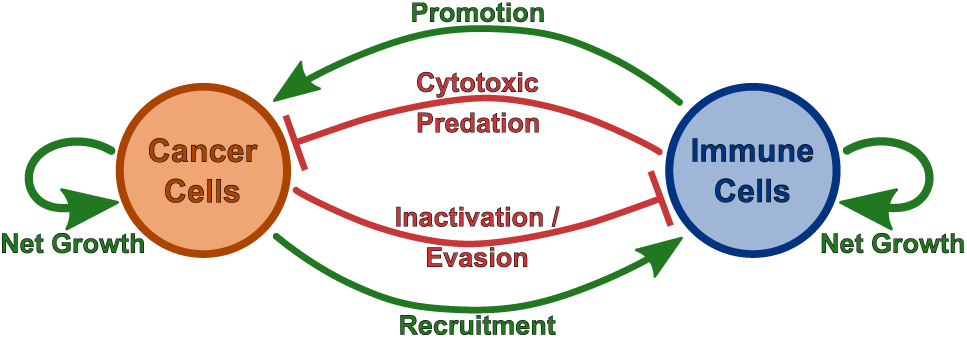
Diagram of cancer-immune interactions that modulate growth dynamics for both cell populations.

In what follows, we prescribe equations to describe the growth of two cell populations over time, cancer cells *C(t)* and immune cells *I(t)*. The main assumption of this model is that the cell populations are homogenous, allowing any spatial dependence to be neglected. Each population is assumed to grow in a self-limited logistic manner with carrying capacities (or maximum possible population sizes) *K*_*C*_ for cancer and *K*_*I*_ for immune.

### A. The Governing Equations

A simple translation of cancer-immune interactions into mathematical form could be along the lines of: the rate of change of the cancer population consists of a net growth term, less an immune-induced predation term, plus an immune-induced stimulation term. Such an additive formulation, however, does not account for the manner in which predation and stimulation occur - which can be tied directly to the ability of cells to sense signals from their environment and proliferate, quiesce, or die accordingly. We thus propose an integrated approach that allows immune actions to modify the net growth term through the following form:

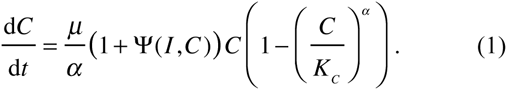

Here μ and α are parameters that describe the growth and sensitivity of cancer cells to environmental regulatory signals, and Ψ is a function that can describe both immune inhibition or stimulation of cancer growth. An example of a predatory / inhibitory functional form is

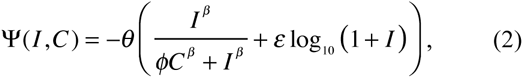

where *θ, β, ϕ*, and *ε* are immune predation parameters.

The immune population is also described by a modified logistic growth formulation,

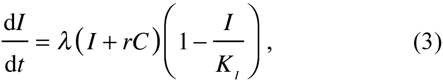

where *λ* is the growth rate and *r* is a recruitment parameter.

An extension of this model allows the cancer to modify its own carrying capacity via [9], [29]

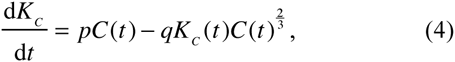

where *p* and *q* are growth stimulation and inhibition constants, respectively. We may further extend this model to allow carrying capacities to depend on both populations [10], (capturing, for example, immune stimulation through inflammation and angiogenesis) and generally describe them by 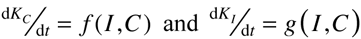.

Parameter values should be determined by fitting the model to experimental or clinical data. Sensitivity analyses can then be used to identify significant determinants of the dynamics in question and ‘prune’ the formalism of less informative parameters.

### B. When this method is useful

We have described how ODEs can be used to track the evolution over time of two populations of cells as they grow within a host. These methods can be used to describe any time evolution, ranging from cellular mass to protein concentration and beyond. Using ODEs to model a biological question requires that there is only one independent variable (such as time) and that all others (such as space) can be neglected. In this case, the structure can simplify over the alternative partial differential equation framework that would be needed if space and time were important, and provide rapid and efficient insights into how the complex immune-cancer dynamic might respond in the clinical context.

## ACKNOWLEDGEMENT

We thank the ICBP and organizers of the joint ICBP-PSOC Mathematical Modelers Meeting (Tampa FL, 2015) for putting together this handbook and for the opportunity to contribute.

